## Supplementary Information for "MME^+^ fibro-adipogenic progenitors are the dominant adipogenic population during fatty infiltration in human skeletal muscle"

A

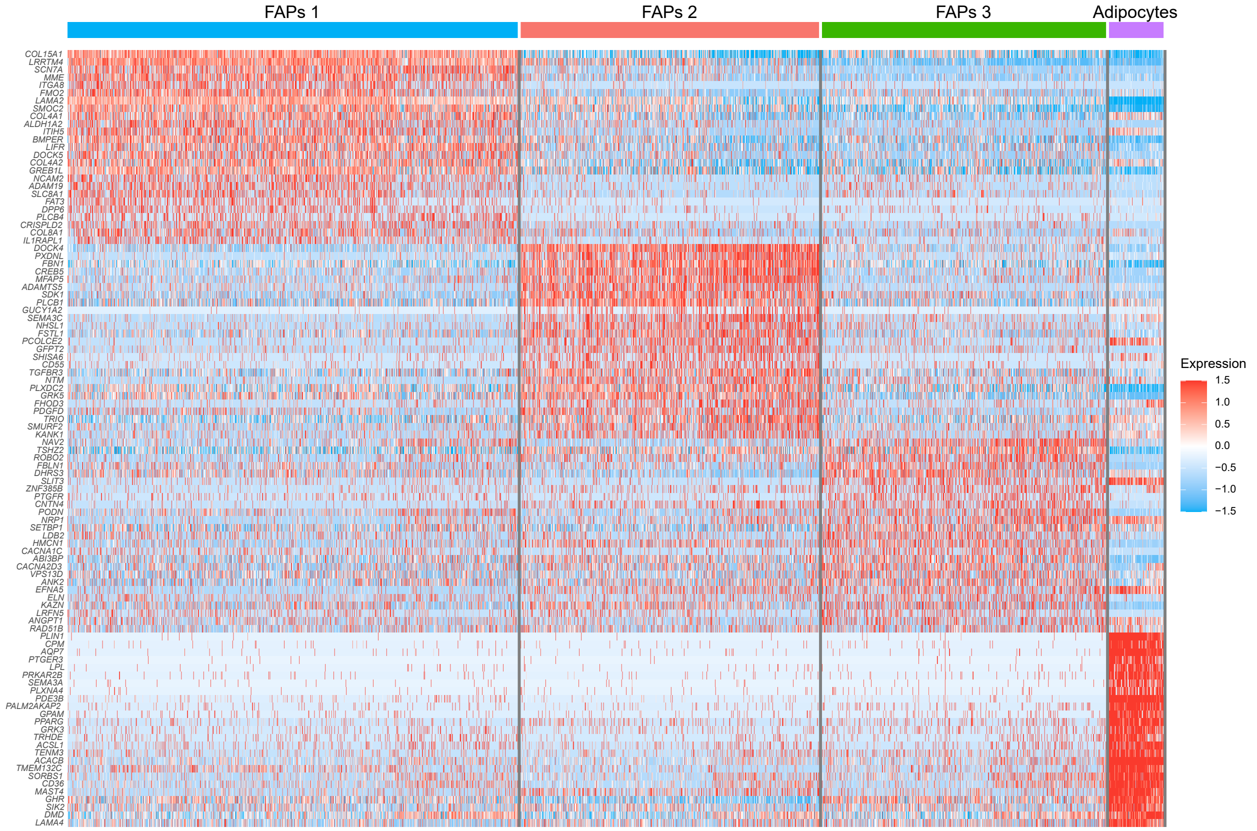

FAPs 1
FAPs 2
FAPs 3
Adipocytes
COL15A1
LRRTM4
SCN7A
MME
ITGA8
FMO2
LAMA2
SMOC2
COL4A1
ALDH1A2
ITIH5
BMPER
LIFR
DOCK5
COL4A2
GREB1L
NCAM2
ADAM19
SLC8A1
FAT3
DPP6
PLCB4
CRISPLD2
COL8A1
IL1RAPL1
DOCK4
PXDNL
FBN1
CREB5
MFAP5
ADAMTS5
SDK1
PLCB1
GUCY1A2
SEMA3C
NHSL1
FSTL1
PCOLCE2
GFPT2
SHISA6
CD55
TGFBR3
NTM
PLXDC2
GRK5
FHOD3
PDGFD
TRIO
SMURF2
KANK1
NAV2
TSHZ2
ROBO2
FBLN1
DHRS3
SLIT3
ZNF385B
PTGFR
CNTN4
PODN
NRP1
SETBP1
LDB2
HMCN1
CACNA1C
ABI3BP
CACNA2D3
VPS13D
ANK2
EFNA5
ELN
KAZN
LRFN5
ANGPT1
RAD51B
PLIN1
CPM
AQP7
PTGER3
LPL
PRKAR2B
SEMA3A
PLXNA4
PDE3B
PALM2AKAP2
GPAM
PPARG
GRK3
TRHDE
ACSL1
TENM3
ACACB
TMEM132C
SORBS1
CD36
MAST4
GHR
SIK2
DMD
LAMA4
Expression
1.5
1.0
0.5
0.0
−0.5
−1.0
−1.5


B

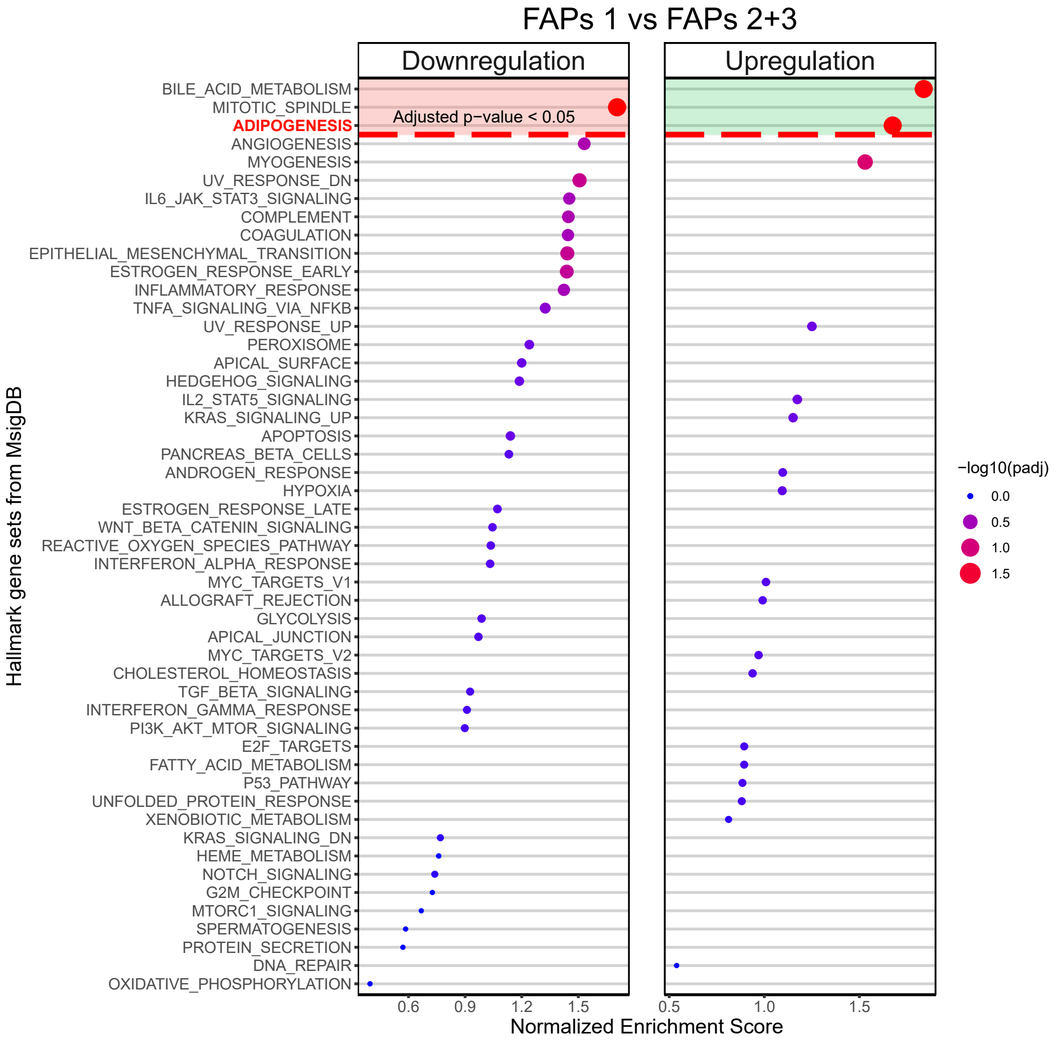

FAPs 1 vs FAPs 2+3
Downregulation
Upregulation
Adjusted p−value < 0.05
Hallmark gene sets from MsigDB
BILE_ACID_METABOLISM
MITOTIC_SPINDLE
ADIPOGENESIS
ANGIOGENESIS
MYOGENESIS
UV_RESPONSE_DN
IL6_JAK_STAT3_SIGNALING
COMPLEMENT
COAGULATION
EPITHELIAL_MESENCHYMAL_TRANSITION
ESTROGEN_RESPONSE_EARLY
INFLAMMATORY_RESPONSE
TNFA_SIGNALING_VIA_NFKB
UV_RESPONSE_UP
PEROXISOME
APICAL_SURFACE
HEDGEHOG_SIGNALING
IL2_STAT5_SIGNALING
KRAS_SIGNALING_UP
APOPTOSIS
PANCREAS_BETA_CELLS
ANDROGEN_RESPONSE
HYPOXIA
ESTROGEN_RESPONSE_LATE
WNT_BETA_CATENIN_SIGNALING
REACTIVE_OXYGEN_SPECIES_PATHWAY
INTERFERON_ALPHA_RESPONSE
MYC_TARGETS_V1
ALLOGRAFT_REJECTION
GLYCOLYSIS
APICAL_JUNCTION
MYC_TARGETS_V2
CHOLESTEROL_HOMEOSTASIS
TGF_BETA_SIGNALING
INTERFERON_GAMMA_RESPONSE
PI3K_AKT_MTOR_SIGNALING
E2F_TARGETS
FATTY_ACID_METABOLISM
P53_PATHWAY
UNFOLDED_PROTEIN_RESPONSE
XENOBIOTIC_METABOLISM
KRAS_SIGNALING_DN
HEME_METABOLISM
NOTCH_SIGNALING
G2M_CHECKPOINT
MTORC1_SIGNALING
SPERMATOGENESIS
PROTEIN_SECRETION
DNA_REPAIR
OXIDATIVE_PHOSPHORYLATION
0.6
0.9
1.2
1.5
0.5
1.0
1.5
Normalized Enrichment Score
−log10(padj)
0.0
0.5
1.0
1.5

**Figure S1. snRNA-seq analysis from human muscle identifies 3 FAPs populations. Related to Figure 1. A)** Heatmap with Z scores showing top enriched marker genes per cell type. Each row represents a gene and each column a cell. Columns are ordered and color-coded based on the cluster of origin. **B)** Dot plots showing downregulated (left panel) or upregulated (right panel) pathways from GSEA of FAPs 1 compared to FAPs 2+3. X-axis indicates the normalized enrichment score (NES) for each pathway. Color and size of the dots indicate adjusted p-values (FDR). Significant values are delimited by the red dashed line (Adjusted p-value < 0.05 or -log10(adjusted p-value) > 1.3).

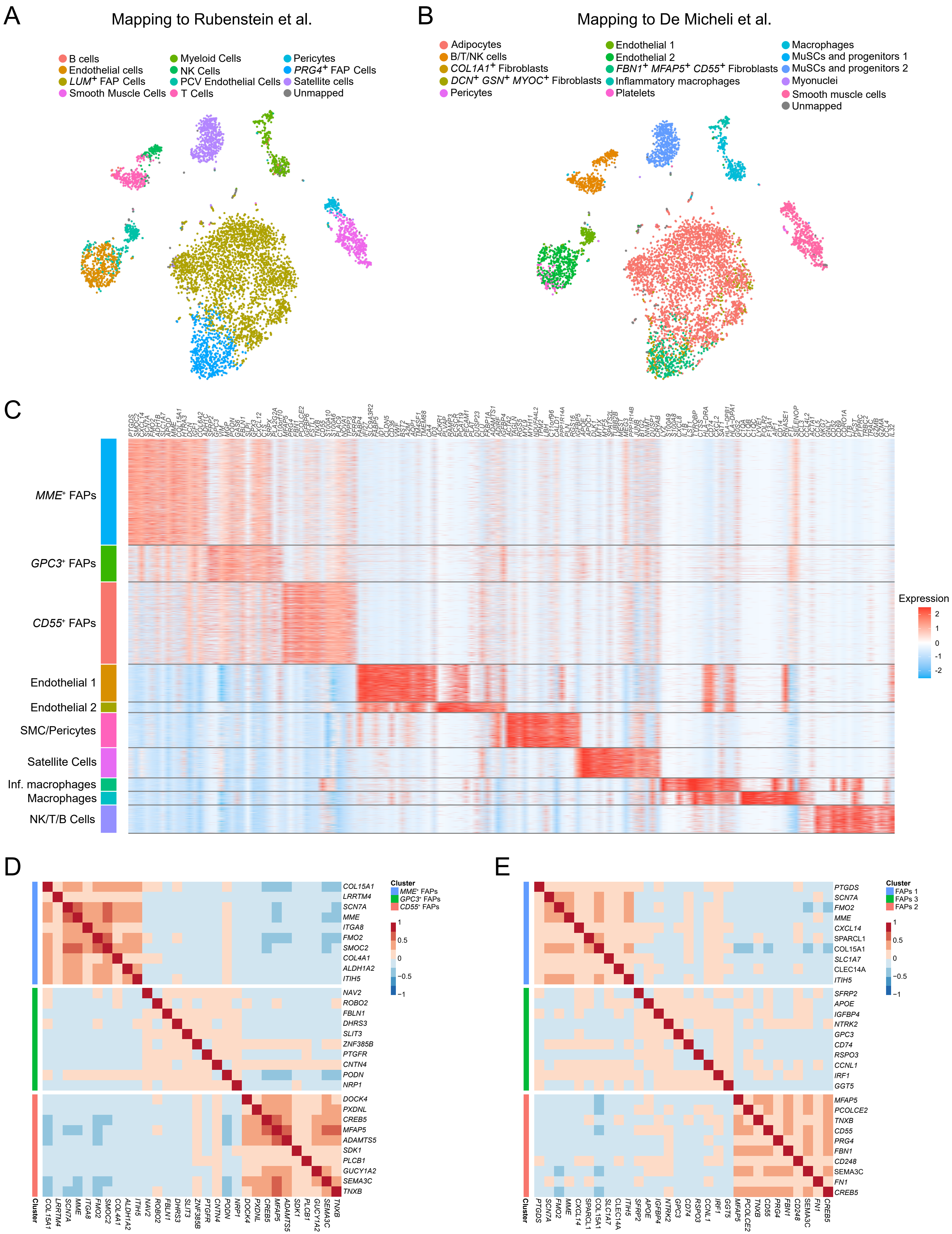

A
Mapping to Rubenstein et al.
B cells
Endothelial cells
LUM+ FAP Cells
Smooth Muscle Cells
Myeloid Cells
NK Cells
PCV Endothelial Cells
T Cells
Pericytes
PRG4+ FAP Cells
Satellite cells
Unmapped
B
Mapping to De Micheli et al.
Adipocytes
B/T/NK cells
COL1A1+ Fibroblasts
DCN+ GSN+ MYOC+ Fibroblasts
Pericytes
Endothelial 1
Endothelial 2
FBN1+ MFAP5+ CD55+ Fibroblasts
Inflammatory macrophages
Platelets
Macrophages
MuSCs and progenitors 1
MuSCs and progenitors 2
Myonuclei
Smooth muscle cells
Unmapped
C
MME+ FAPs
GPC3+ FAPs
CD55+ FAPs
Endothelial 1
Endothelial 2
SMC/Pericytes
Satellite Cells
Inf. macrophages
Macrophages
NK/T/B Cells
Expression
2
1
0
-1
-2
D
COL15A1
LRRTM4
SCN7A
MME
ITGA8
FMO2
SMOC2
COL4A1
ALDH1A2
ITIH5
NAV2
ROBO2
FBLN1
DHRS3
SLIT3
ZNF385B
PTGFR
CNTN4
PODN
NRP1
DOCK4
PXDNL
CREB5
MFAP5
ADAMTS5
SDK1
PLCB1
GUCY1A2
SEMA3C
TNXB
Cluster
MME+ FAPs
GPC3+ FAPs
CD55+ FAPs
1
0.5
0
−0.5
−1
E
PTGDS
SCN7A
FMO2
MME
CXCL14
SPARCL1
COL15A1
SLC1A7
CLEC14A
ITIH5
SFRP2
APOE
IGFBP4
NTRK2
GPC3
CD74
RSPO3
CCNL1
IRF1
GGT5
MFAP5
PCOLCE2
TNXB
CD55
PRG4
FBN1
CD248
SEMA3C
FN1
CREB5
Cluster
FAPs 1
FAPs 3
FAPs 2
1
0.5
0
−0.5
−1

**Figure S2. scRNA-seq analysis from human muscle identifies main FAPs and mononuclear cell populations. Related to Figure 2. A)** TSNE plot of main cell populations identified in human muscle by the automatic mapping to Rubenstein et al. dataset, color-coded by the identified populations. **B)** Same as A) but mapping to De Micheli et al. dataset. **C)** Heatmap with Z scores showing top enriched marker genes per cell type. Each column represents a gene and each row a cell. Rows are ordered and color-coded based on the cluster of origin. **D)** Heatmap representing Pearson's correlation values of human FAP markers from the snRNA-seq dataset in the human FAPs scRNA-seq dataset. Horizontal color-coded separation indicates the set of marker genes from each FAP subpopulation. **E)** Heatmap representing Pearson's correlation values of human FAP markers from the scRNA-seq dataset in the human FAPs snRNA-seq dataset. Horizontal color-coded separation indicates the set of marker genes from each FAP subpopulation.

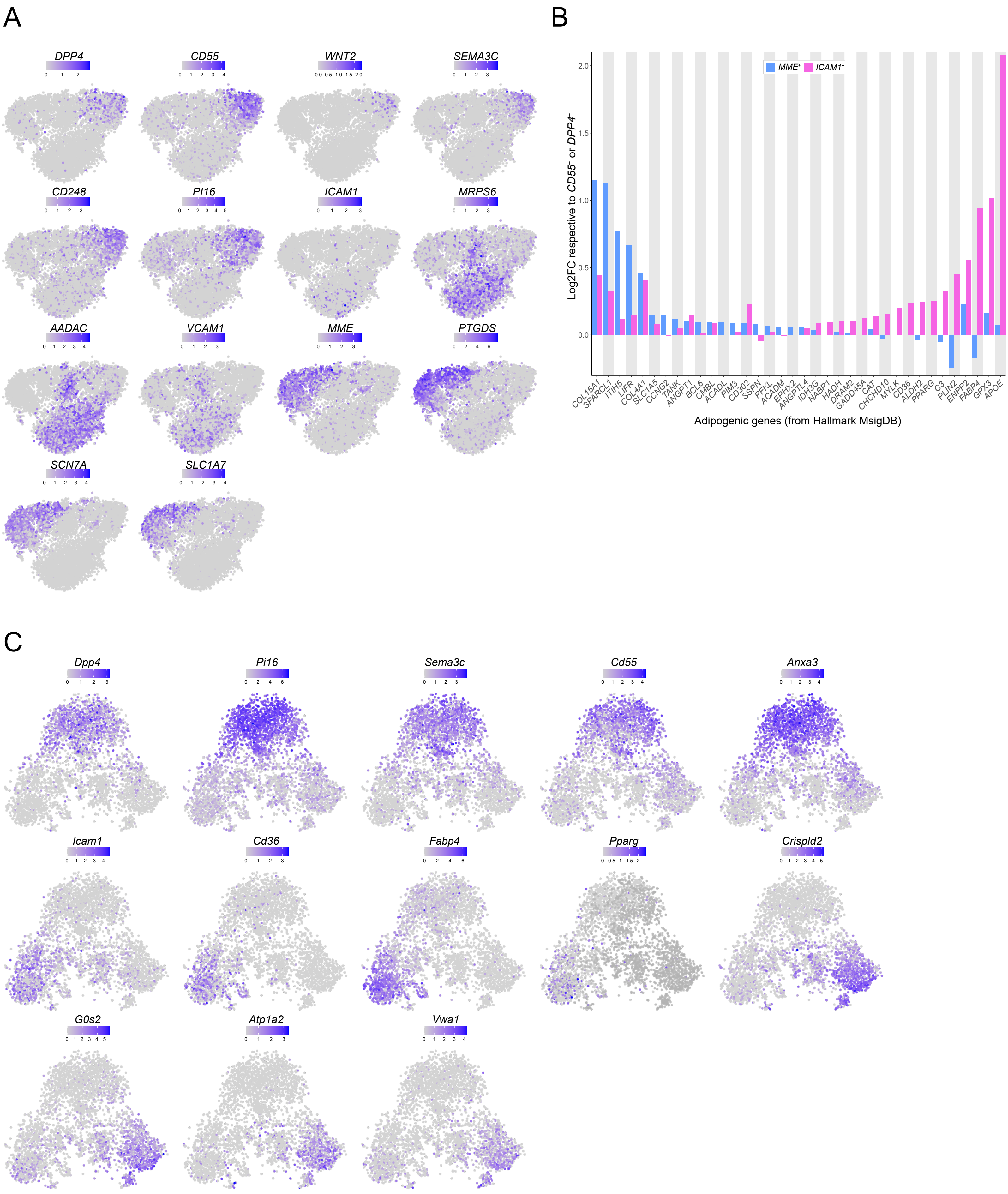
A
DPP4
CD55
WNT2
SEMA3C
CD248
PI16
ICAM1
MRPS6
AADAC
VCAM1
MME
PTGDS
SCN7A
SLC1A7
B
MME⁺
ICAM1⁺
Log2FC respective to CD55⁺ or DPP4⁺
COL15A1
SPARCL1
ITIH5
LIFR
COL4A1
SLC1A5
CCNG2
TANK
ANGPT1
BCL6
CMBL
ACADL
PIM3
CD302
SSPN
PFKL
ACADM
EPHX2
ANGPTL4
IDH3G
NABP1
HADH
DRAM2
GADD45A
CAT
CHCHD10
MYLK
CD36
ALDH2
PPARG
C3
PLIN2
ENPP2
FABP4
GPX3
APOE
Adipogenic genes (from Hallmark MsigDB)
C
Dpp4
Pi16
Sema3c
Cd55
Anxa3
Icam1
Cd36
Fabp4
Pparg
Crispld2
G0s2
Atp1a2
Vwa1

**Figure S3. *MME*⁺ and *GPC3*⁺ muscle FAPs are not found in adipose tissue. Related to Figure 3. A)** TSNE plots of FAPs marker genes identified in the integration of human muscle FAPs and human adipose progenitor cells from Merrick et al., color-coded by logcounts values. **B)** Log2-FoldChange of adipogenic genes (from Hallmark MsigDB) found to be upregulated in *MME*$^+$ or *ICAM1*$^+$ cells. Each log2-FoldChange is relative to *CD55*$^+$ or *DPP4*$^+$ for *MME*$^+$ and *ICAM1*$^+$ cells respectively. **C)** Same as A) but FAPs marker genes identified in the integration of mouse muscle FAPs and adult mouse (10 weeks) adipose progenitor cells from Merrick et al. color-coded by logcounts values.

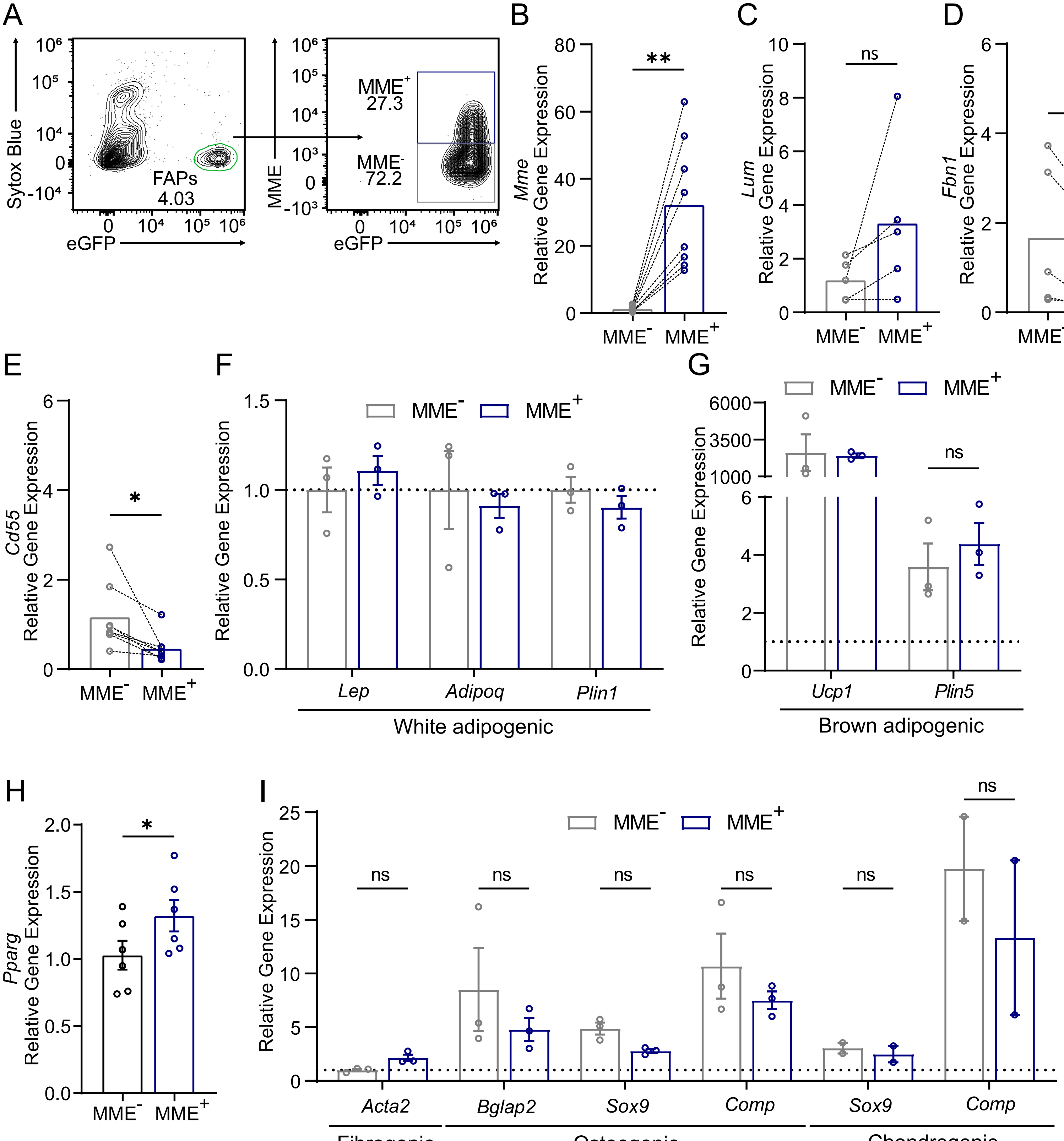

A
Sytox Blue
eGFP
FAPs
4.03
MME
MME+
27.3
MME-
72.2
B
Mme
Relative Gene Expression
**
MME-
MME+
C
Lum
Relative Gene Expression
ns
D
Fbn1
Relative Gene Expression
*
E
Cd55
Relative Gene Expression
*
F
Relative Gene Expression
Lep
Adipoq
Plin1
White adipogenic
G
Relative Gene Expression
ns
Ucp1
Plin5
Brown adipogenic
H
Pparg
Relative Gene Expression
*
I
Relative Gene Expression
ns
Acta2
Bglap2
Sox9
Comp
Sox9
Comp
Fibrogenic
Osteogenic
Chondrogenic

**Figure S4. MME$^+$ are a highly adipogenic fraction of PDGFR$\alpha^+$ FAPs. Related to Figure 4. A)** Representative FACS plot outlining the sorting strategy to isolate PDGFR$\alpha$-eGFP$^+$MME$^{-/+}$ FAPs. **B)** Relative expression of *Mme* in freshly sorted MME$^{-/+}$ FAPs. **C)** Relative expression of *Lum* in freshly sorted MME$^{-/+}$ FAPs. **D)** Relative expression of *Fbn1* in freshly sorted MME$^{-/+}$ FAPs. **E)** Relative expression of *Cd55* in freshly sorted MME$^{-/+}$ FAPs. **F)** Relative gene expression of *Lep, Adipoq,* and *Plin1* in MME$^{-/+}$ FAPs derived adipocytes differentiated in full white adipogenic medium. Related to Figure 4E. **G)** Relative gene expression of *Ucp1* and *Plin5* in MME$^{-/+}$ FAPs derived brown adipocytes normalized to their expression in MME$^-$ FAPs derived white adipocytes (dotted line). **H)** Relative gene expression of *Pparg* in MME$^{-/+}$ FAPs derived adipocytes differentiated in low insulin medium. **I)** Relative gene expression of *Acta2, Bglap2, Sox9, and Comp* in MME$^{-/+}$ FAPs derived fibroblasts, osteocytes, or chondrocytes as indicated. The expression of all genes was normalized to their expression in MME$^-$ FAPs derived white adipocytes (dotted line). Each dot represents a single mouse. Bar graphs represent the mean in **B**-**E** and represent the mean ± SEM in **F**-**I**. Student's t-test (two tailed, paired, *$p<0.05$, **$p<0.01$, ns>0.05) was used in **B**-**E** and **H**. Two-way ANOVA with Šidáks multiple comparison test (ns>0.05) was used in **F**, **G**, and **I**.

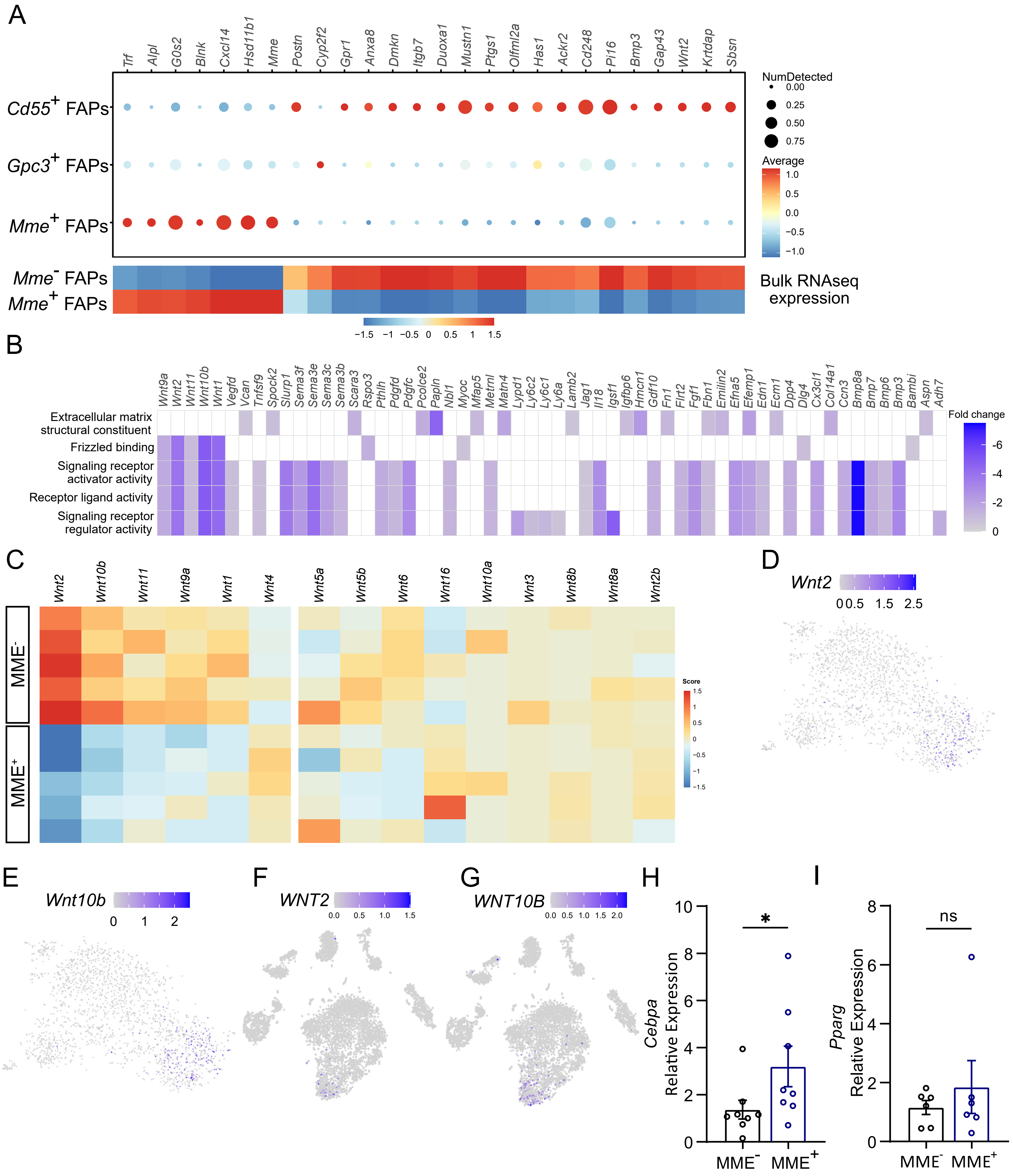

A
Trf Alpl G0s2 Blnk Cxcl14 Hsd11b1 Mme Postn Cyp2f2 Gpr1 Anxa8 Dmkn Itgb7 Duoxa1 Mustn1 Ptgs1 Olfml2a Has1 Ackr2 Cd248 Pi16 Bmp3 Gap43 Wnt2 Krtdap Sbsn
Cd55+ FAPs
Gpc3+ FAPs
Mme+ FAPs
NumDetected
0.00
0.25
0.50
0.75
Average
1.0
0.5
0.0
−0.5
−1.0
Mme- FAPs
Mme+ FAPs
Bulk RNAseq expression
−1.5 −1 −0.5 0 0.5 1 1.5
B
Wnt9a Wnt2 Wnt11 Wnt10b Wnt1 Vegfd Vcan Tnfsf9 Spock2 Slurp1 Sema3f Sema3e Sema3c Sema3b Scara3 Rspo3 Pthlh Pdgfd Pdgfc Pcolce2 Papln Nbl1 Myoc Mfap5 Metrnl Matn4 Lypd1 Ly6c2 Ly6c1 Ly6a Lamb2 Jag1 Il18 Igsf1 Igfbp6 Hmcn1 Gdf10 Fn1 Flrt2 Fgf1 Fbn1 Emilin2 Efna5 Efemp1 Edn1 Ecm1 Dpp4 Dlg4 Cx3cl1 Col14a1 Ccn3 Bmp8a Bmp7 Bmp6 Bmp3 Bambi Aspn Adh7
Extracellular matrix structural constituent
Frizzled binding
Signaling receptor activator activity
Receptor ligand activity
Signaling receptor regulator activity
Fold change
-6
-4
-2
0
C
Wnt2 Wnt10b Wnt11 Wnt9a Wnt1 Wnt4 Wnt5a Wnt5b Wnt6 Wnt16 Wnt10a Wnt3 Wnt8b Wnt8a Wnt2b
MME-
MME+
Score
1.5
1
0.5
0
−0.5
−1
−1.5
D
Wnt2
0 0.5 1.5 2.5
E
Wnt10b
0 1 2
F
WNT2
0.0 0.5 1.0 1.5
G
WNT10B
0.0 0.5 1.0 1.5 2.0
H
Cebpa Relative Expression
0 2 4 6 8 10
*
MME- MME+
I
Pparg Relative Expression
0 2 4 6 8
ns
MME- MME+

**Figure S5. MME$^+$ FAPs are characterized by reduced WNT signaling and refractory to WNT-mediated inhibition of adipogenesis. Related to Figure 5. A)** Dot plot from mouse scRNA-seq of high variable genes identified in the mouse bulk RNA-seq (see figure 5B). Each row represents a FAP subpopulation and each column a gene. Color and size of the dots indicate the level of expression and the percentage of cells that express the gene respectively. The simplified heatmap below displays Z-scores averaged per cell type from the bulk RNA-seq data set (see Figure 5B). **B)** Heatmap showing the top 5 significantly downregulated *molecular function* processes (adjusted p-value < 0.05) in a Gene Ontology (GO) enrichment analysis in *MME*$^+$ FAPs from the mouse bulk RNA-seq. Color-coded by the log2-FoldChange respective to *MME*$^-$ FAPs. Each row represents a *molecular function* process and each column a gene which is significantly regulated within our dataset that belongs to that *molecular function* process (log2-FoldChange > 0.5 and FDR < 0.05). Blank tiles indicate that a gene does not belong to that *molecular function* process. **C)** Heatmap with Z scores showing WNT ligands in MME$^{-/+}$ FAPs. Genes displayed left of the line break are differentially expressed (log2-FoldChange > 0.5 and FDR < 0.05). **D)** TSNE plot of *Wnt2* color-coded by logcounts values in mouse FAPs **E)** TSNE plot of *Wnt10b* color-coded by logcounts values in mouse FAPs **F)** TSNE plot of *WNT2* color-coded by logcounts values in FAPs in human muscle. **G)** TSNE plot of *WNT10B* color-coded by logcounts values in FAPs in human muscle. **H-I)** Relative expression of *Cebpa* (**H**) and *Pparg* (**I**) in freshly sorted MME$^{-/+}$ FAPs. Each dot represents a single mouse. Bar graphs represent the mean ± SEM. Student's t-test (two tailed, paired, ns>0.05, *p<0.05) was used in **H** and **I**.

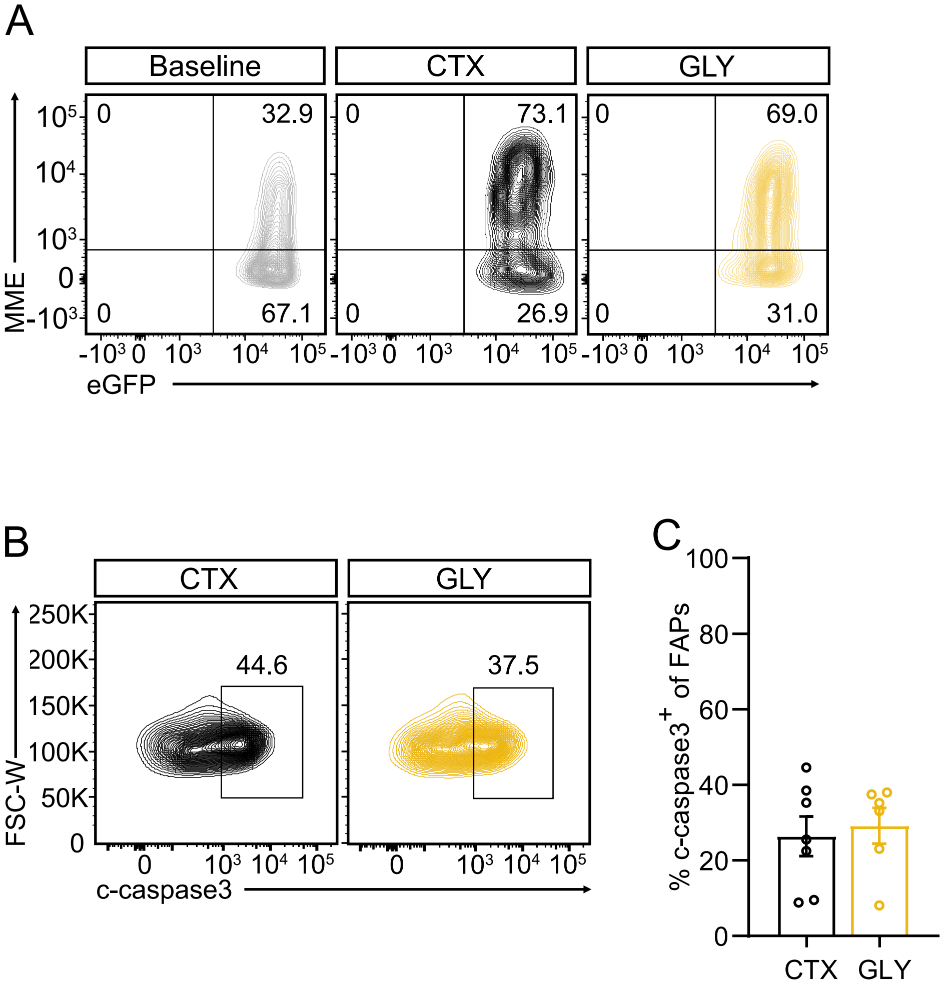
A
Baseline
CTX
GLY
MME
eGFP
32.9
67.1
73.1
26.9
69.0
31.0
B
CTX
GLY
FSC-W
c-caspase3
44.6
37.5
C
% c-caspase3+ of FAPs
CTX
GLY

**Figure S6. MME$^+$ FAPS are more prone to apoptosis after injury and deplete upon glycerol injury. Related to Figure 6. A)** Representative flow cytometric analysis of eGFP$^+$ FAPs in CTX and GLY injected muscle at 3 days post injury (dpi) **B-C)** Representative flow cytometric analysis (**B**) and quantification (**C**) of c-caspase3$^+$ FAPs as a percentage of eGFP$^+$ FAPs at 4 dpi in CTX injected and GLY injected muscle. Each dot represents a single mouse. Bar graphs represent the mean ± SEM. Student's t-test (two tailed, unpaired, **$p<0.01$) was used in **C**.

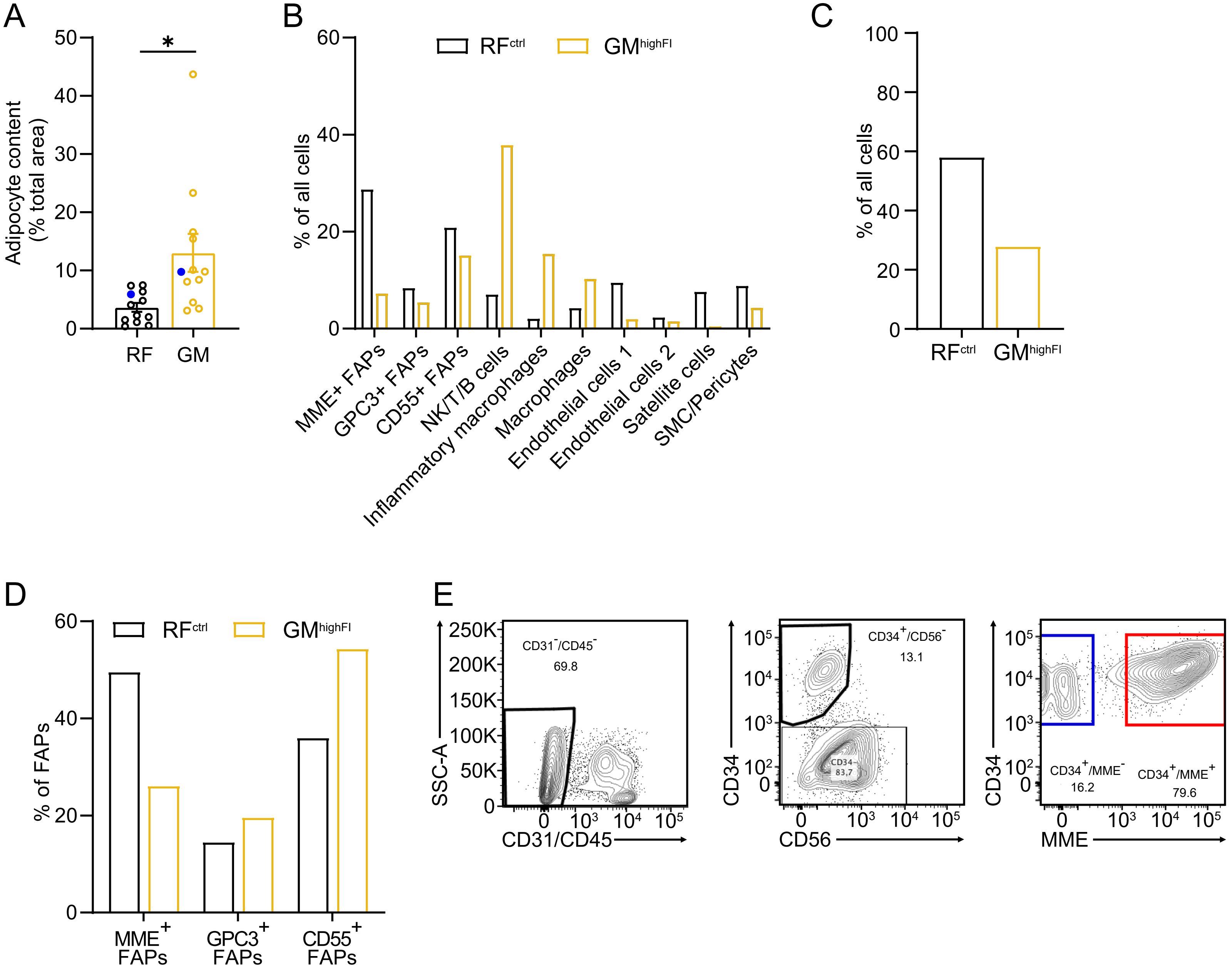
A
Adipocyte content (% total area)
*
RF
GM
B
% of all cells
RFctrl
GMhighFI
MME+ FAPs
GPC3+ FAPs
CD55+ FAPs
NK/T/B cells
Inflammatory macrophages
Macrophages
Endothelial cells 1
Endothelial cells 2
Satellite cells
SMC/Pericytes
C
% of all cells
RFctrl
GMhighFI
D
% of FAPs
RFctrl
GMhighFI
MME+ FAPs
GPC3+ FAPs
CD55+ FAPs
E
SSC-A
CD31/CD45
CD31-/CD45-
69.8
CD34
CD56
CD34+/CD56-
13.1
CD34-
83.7
CD34
MME
CD34+/MME-
16.2
CD34+/MME+
79.6

**Figure S7. Human MME$^+$ FAPs are highly adipogenic and are exhausted in fatty infiltrated human muscle. Related to Figure 7. A)** Relative adipocyte area as quantified by adipocyte area relative to total biopsy area on H&E-stained sections. Biopsies from the patient used for scRNA-seq are marked with blue dots. **B)** Bar plots showing the percentage of the main cell types within each dataset from either control muscle (RF$^{ctrl}$) or highly fatty infiltrated (GM$^{highFI}$) human muscle. **C)** Bar plots showing the percentage of the combined FAP populations within each dataset from either control muscle (RF$^{ctrl}$) or highly fatty infiltrated (GM$^{highFI}$) human muscle. **D)** Bar plots showing the percentage of FAPs relative to the total FAP fraction within each dataset from either control muscle (RF$^{ctrl}$) or highly fatty infiltrated (GM$^{highFI}$) human muscle. **E)** Representative FACs plots outlining the sorting strategy to isolate MME$^{-/+}$ human FAPs. Each dot represents an individual patient. Bar graphs represent mean ± SEM in **A**. Student's t test (two tailed, paired, *p<0.05) was used in **A**. Bar plots in **B, C,** and **D** represent the percentage of all cells or the percentage of FAPs in the RF$^{ctrl}$ and GM$^{highFI}$ single cell datasets.

### Supplemental table

**Table S1. Sequence of primers used for RT-PCR. Related to Figure 5 and Supplementary Figures S4 and S5**

| Gene Name | Forward Primer | Reverse Primer |
|---|---|---|
| *Rna18s5* | AGTCCCTGCCCTTTGTACACA | CGATCCGAGGGCCTCACTA |
| *Wnt2* | ACAACAGAGCTGGAAGGAAGGCTGT | AGTGAAGCCAGTGCCATCCTGG |
| *Wnt10b* | GCTGACTGACTCGCCCACCG | AAGCACACGGTGTTGGCCGT |
| *Mme* | GGGAGGCTTTATGTGGAAGC | CCGGATTTGTGCAATCAAGT |
| *Lum* | CCCACCCTGACAGAGTTCAC | ATTGGCCACTGACACTACCG |
| *Fbn1* | TGAGAGTCCGAGCCGCTAGT | ACAGCTTTCTTCTCCAGGGAC |
| *Cd55* | GGAGAGCCTAACACAGGTGG | TCTTCGTAACTCTTCGTTGGCT |
| *Lep* | TCAAGACCATTGTCACCAGG | TGAAGCCCAGGAATGAAGTC |
| *Adipoq* | TGTTCCTCTTAATCCTGCCCA | CCAACCTGCACAAGTTCCCTT |
| *Ucp1* | GGACGACCCCTAATCTAATGAG | GCAAAACCCGGCAACAAG |
| *Plin5* | CCATCTCGCCTATGAACACTCTT | CAGCTGGGCCAGCATCTC |
| *Plin1* | CTGTGTGCAATGCCTATGAGA | CTGGAGGGTATTGAAGAGCCG |
| *Pparg* | TCGCTGATGCACTGCCTATG | GAGAGGTCCACAGAGCTGATT |
| *Acta2* | GTCCCAGACATCAGGGAGTAA | TCGGATACTTCAGCGTCAGGA |
| *Bglap2* | CTGACCTCACAGATCCCAAGC | TGGTCTGATAGCTCGTCACAAG |
| *Sox9* | AGTACCCGCATCTGCACAAC | ACGAAGGGTCTCTTCTCGCT |
| *Comp* | ACTGCCTGCGTTCTAGTGC | CGCCGCATTAGTCTCCTGAA |
| *Cebpa* | GCCGAGATAAAGCCAAACAAC | GACCCGAAACCATCCTCTG |
